# Interorganelle lipid flux revealed by enzymatic mass tagging *in vivo*

**DOI:** 10.1101/2021.08.27.457935

**Authors:** Arun T. John Peter, Matthias Peter, Benoît Kornmann

## Abstract

The distinct activities of cellular organelles are dependent on the proper function of their membranes. Coordinated membrane biogenesis of the different organelles necessitates interorganelle transport of lipids from their site of synthesis to their destination membranes. Several proteins and trafficking pathways have been proposed to participate in lipid distribution, but despite the basic importance of this process, *in vivo* evidence linking the absence of putative transport pathways to specific transport defects remains scarce. An obvious reason for this scarcity is the near absence of *in vivo* lipid trafficking assays. Here we introduce a versatile method named METALIC (<u>M</u>ass tagging-<u>E</u>nabled <u>T</u>r<u>A</u>cking of <u>L</u>ipids <u>I</u>n <u>C</u>ells) to track interorganelle lipid flux inside living cells. In this strategy, two enzymes, one directed to a “donor” and the other to an “acceptor” organelle, add two distinct mass tags to lipids. Mass spectrometry-based detection of lipids bearing the two mass tags is then used as a proxy for lipid exchange between the two organelles. By applying this approach to ER and mitochondria, we show that the ERMES and Vps13-Mcp1 complexes have lipid transport activity *in vivo*, and unravel their relative contributions to ER-mitochondria lipid exchange.

## Introduction

Organelle function depends on the lipids that constitute their membranes. Indeed, membrane lipids are not only involved as structural barriers, but also serve in the recruitment of specific proteins and storage of energy. Because lipid biosynthesis in eukaryotic cells mostly happens in the endoplasmic reticulum (ER), lipids must be transported from their place of synthesis to all other cellular membranes. Lipid transport was long thought to be a by-product of protein vesicular trafficking, but research over the past decade has shown that cells have evolved mechanisms to mediate the bulk of lipid exchange by non-vesicular means ^1^. Although this mode of transport is associated with all organelles of the endomembrane system, it is especially relevant for organelles like mitochondria that are excluded from vesicular traffic.

Non-vesicular lipid exchange often occurs at membrane contact sites, zones of close apposition (10-30 nm) between organelles, where lipid transport proteins (LTPs) solubilize lipids from the membrane, shield them from the aqueous milieu in a hydrophobic pocket, and catalyze their exchange between the two membranes. In yeast, the ERMES (<u>ER</u>-<u>M</u>itochondria <u>E</u>ncounter <u>S</u>tructure) is a complex of such LTPs implicated in lipid exchange between ER and mitochondria ^2–8^. Three of its four core subunits, namely Mmm1, Mdm12, Mdm34, harbor a lipid-solubilizing SMP (<u>s</u>ynaptotagmin-like <u>m</u>itochondrial lipid-binding <u>p</u>rotein) domain. Surprisingly, however, ERMES deficiency does not seem to prevent ER-mitochondria lipid exchange ^2,9^ though it results in a host of phenotypes including slow growth and defective mitochondrial morphology. Another LTP implicated in mitochondrial lipid transport is the conserved chorein-N domain-containing protein Vps13, which associates with mitochondria via its receptor protein Mcp1 ^10–12^. ERMES inactivation when combined with a *vps13* or *mcp1* deletion leads to a synthetic lethal phenotype ^11,13^, suggesting that Vps13 may partially compensate lipid transport deficiency in the absence of ERMES function. However, although both the ERMES complex and Vps13 exhibit lipid transport activity *in vitro* ^3–5,14^, their redundant involvement in lipid exchange remains to be proven *in vivo*.

Indeed, despite the identification of multiple other LTPs localized to various membrane contact sites, our knowledge on inter-organelle lipid transport and the LTP function *in vivo* remains poor, mainly due to limitations in the existing tools to assay lipid transport. For instance, our understanding on the transport of the major lipid class, phospholipids, is mainly derived from two methods: a) *in vivo* assays using radiolabeled precursors and, b) *in vitro* lipid exchange assays ^14–17^. In typical *in vivo* assays, cells are treated with ^3^H-serine, which results in the production of ^3^H-phosphatidylserine (PS) by the PS-synthesizing enzyme in the endoplasmic reticulum (ER). As the PS decarboxylase Psd1 that produces phosphatidylethanolamine (PE) from PS is localized to the inner mitochondrial membrane (IMM) and the methyltransferases, Cho2 and Opi3, that make phosphatidylcholine (PC) from PE are exclusively localized to the ER, detection of ^3^H-PE and ^3^H-PC serve as a read-out for ER-mitochondria lipid exchange. This assay has several limitations. First, it is limited to the ER and mitochondria. Second, phospholipases might release labeled lipid headgroups, which can be reincorporated into phospholipids via the alternative PE- and PC-synthesizing Kennedy pathway in the ER, independent of interorganelle lipid transport. Finally, the recent finding that a fraction of Psd1 is localized to the ER in addition to the IMM ^18^ weakens the validity of this assay as all the three phospholipids PS, PE and PC can be made in the ER without the need for ER-mitochondria lipid exchange. Although *in vitro* assays that monitor lipid exchange between liposomes ^14,17^ have been very useful to test the sufficiency of LTPs implicated in lipid transport, they do not inform about lipid exchange rates, identity, origin and destinations, and regulation of lipid transport routes *in vivo*.

To address these limitations, we have developed a novel assay called METALIC (<u>M</u>ass tagging-<u>E</u>nabled <u>T</u>r<u>A</u>cking of <u>L</u>ipids <u>I</u>n <u>C</u>ells) in which we exploit enzyme-mediated mass-tagging to measure the exchange of specific lipids between two organelles *in vivo*. Using this versatile approach, we unravel lipid transport activity of Vps13 and the ERMES complex *in vivo*, and quantify their relative contributions in ER-mitochondria lipid exchange.

## Results

### Principle of the METALIC assay

The principle of METALIC is to target a first lipid-modifying enzyme to an organelle of interest where it chemically modifies lipids at that specific “donor” compartment, introducing a diagnostic “mass tag.” Upon transport to an “acceptor” compartment, the mass-tagged lipid encounters a second enzyme that introduces a different “mass tag”. The detection of doubly mass-tagged lipids by mass spectrometry thus serves as a proxy to monitor lipid transport between the two compartments. Importantly, this mass-tagging approach can be combined with metabolic labeling to capture the kinetics of lipid transport. For instance, pulse labeling with deuterated precursors can be used to assess the kinetics of appearance of i) each labeled mass tag separately, a proxy for the activity of each enzyme and the metabolic activity of the cell, and ii) doubly mass-tagged lipids, a proxy for lipid exchange rates between the donor and acceptor organelles.

### The bacterial cyclopropane fatty acid synthase is active and targetable in yeast

We made use of the cyclopropane-fatty-acyl phospholipid synthase (CFAse), a soluble bacterial enzyme that introduces a methylene group (-CH_2_-) at double bonds of fatty acyl chains in phospholipids, using *S*-adenosyl methionine (SAM) as a cofactor (Fig 1A) ^19^. The resulting cyclopropane lipids have similar biophysical properties as their unsaturated precursors ^20^, but carry an identifiable +14-Dalton mass tag (Fig 1A). To test its enzymatic properties in a heterologous environment, we expressed CFAse constitutively in yeast and used mass spectrometry to assay for the modification of the most abundant phospholipid species, phosphatidylcholine (PC). The mass spectrum revealed the appearance of peaks 14 Da heavier than the precursor PC species upon expression of CFAse, confirming that the enzyme is active in yeast (Fig 1B). The modified lipids represented up to 50 % of the amount of any particular precursor species. To test if CFAse can be targeted to specific organelles, we fused various targeting sequences (see Materials and Methods) to a CFAse-mCherry construct and verified proper localization by microscopy (Fig 1C). Expression of CFAse in different organelles did not prevent cell growth, demonstrating that cells tolerated the production of cyclopropane fatty acids at the tested organelles (Fig 1D).

**Figure 1.**
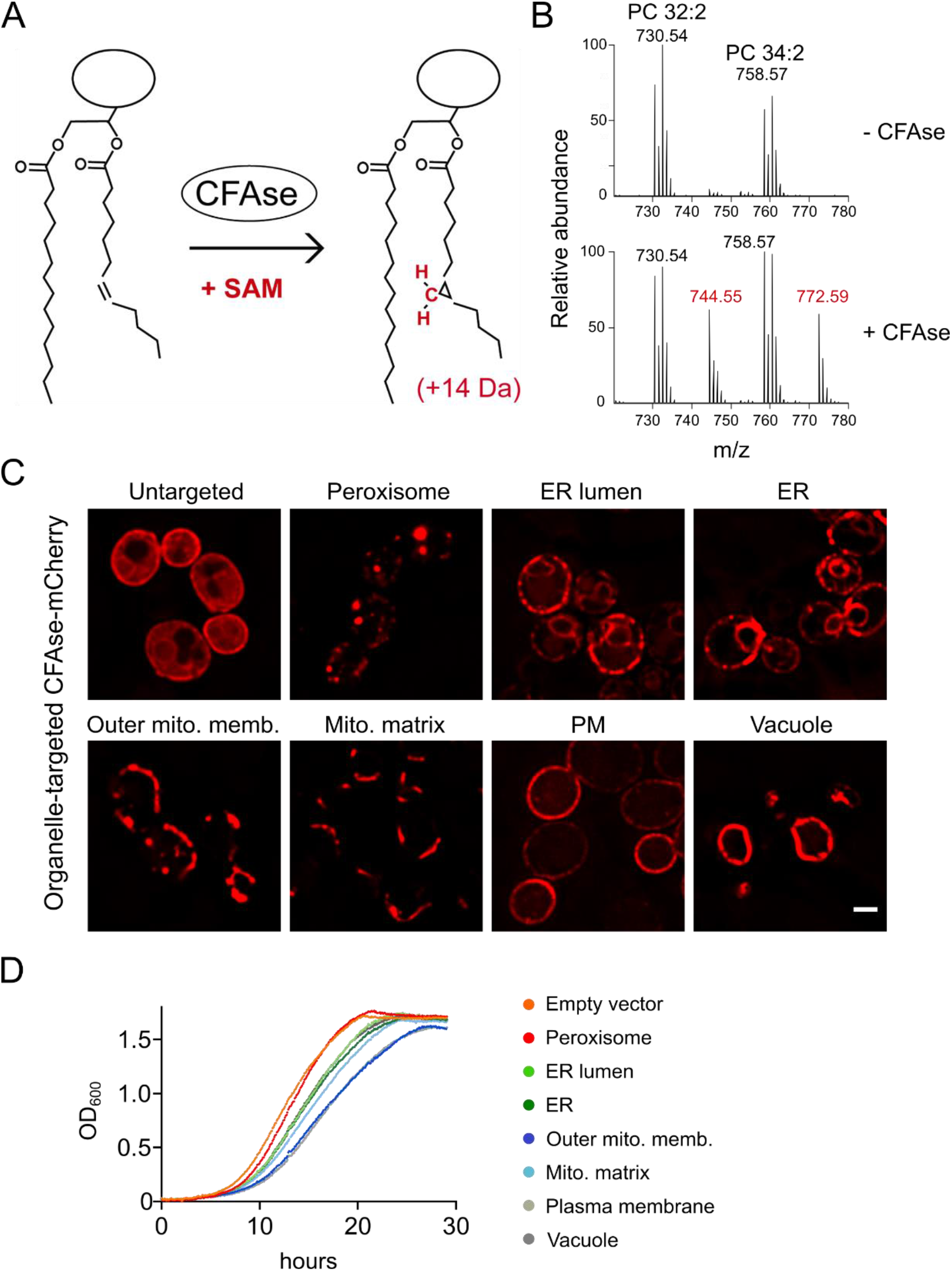
*(A)* Scheme depicting the CFAse reaction. CFAse adds a methylene group to double bonds on fatty acyl chains in phospholipids irrespective of their head group, using *S*-adenosyl methionine (SAM) as a co-factor, resulting in a +14 Dalton mass-shift. *(B)* CFAse is active in yeast. Mass spectrum showing phosphatidylcholine species PC 32:2 (730.54 m/z) and PC 34:2 (758.57 m/z) and their mass-tagging upon expression of CFAse in yeast. *(C)* Localization of mCherry-tagged CFAse to different organelles by fusion to targeting sequences (see Material and Methods). Scale bar, 2 μm. *(D)* Growth of yeast cells expressing mCherry-tagged CFAse constructs targeted to different organelles was monitored by OD_600_ measurements.

#### CFAse mass-tags various phospholipids in different organelles

To assay the activity of CFAse targeted to different organelles, we quantified whole-cell phospholipids including phosphatidylserine (PS), phosphatidylethanolamine (PE), phosphatidylcholine (PC), phosphatidylinositol (PI) and phosphatidylglycerol (PG) in wild-type cells using mass spectrometry. We found that for all the different phospholipids species upon targeting to different organelles, we could detect cyclopropylated (+14 Da) molecules, the percentage of which is shown for any given precursor species (Fig 2). Both distinct as well as similar patterns in the extent of mass-tagging of different phospholipids were detected depending upon the targeted organelle. As a striking example, in cells expressing the mitochondria matrix-targeted CFAse, PG and PE represented the highest mass-tagged fraction relative to other phospholipid species, in line with these two lipids being synthesized and accumulating in the mitochondrial inner membrane. In most cases, modified lipids originating from precursors with two double bonds were more abundant than those originating from lipids with a single double bond, consistent with the idea that phospholipids with two unsaturated fatty acids double the chance for CFAse modification, and indicating that CFAse might have little preference for the one or the other fatty acid (*sn*-1 or *sn*-2). Interestingly, the relative abundance of the mass-tagged species originating from PE 32:2, 32:1 and 34:2 was comparable in all targeted destinations except for mitochondrial matrix where tagging of the mono-unsaturated 32:1 species dominated over the 32:2 and 34:2 species. On the other hand, in the case of PC, the relative abundance profile followed the order 32:2>34:2>32:1>34:1 irrespective of the organelle to which CFAse was targeted, a pattern that mimics the abundance profile observed in the whole cell lipidome of yeast cells (20). Therefore, differences in modification efficiency can be explained by differences in substrate abundance. Taken together, these results demonstrate that the bacterial CFAse can be used specifically and efficiently tag phospholipids in distinct organelles.

**Figure 2.**
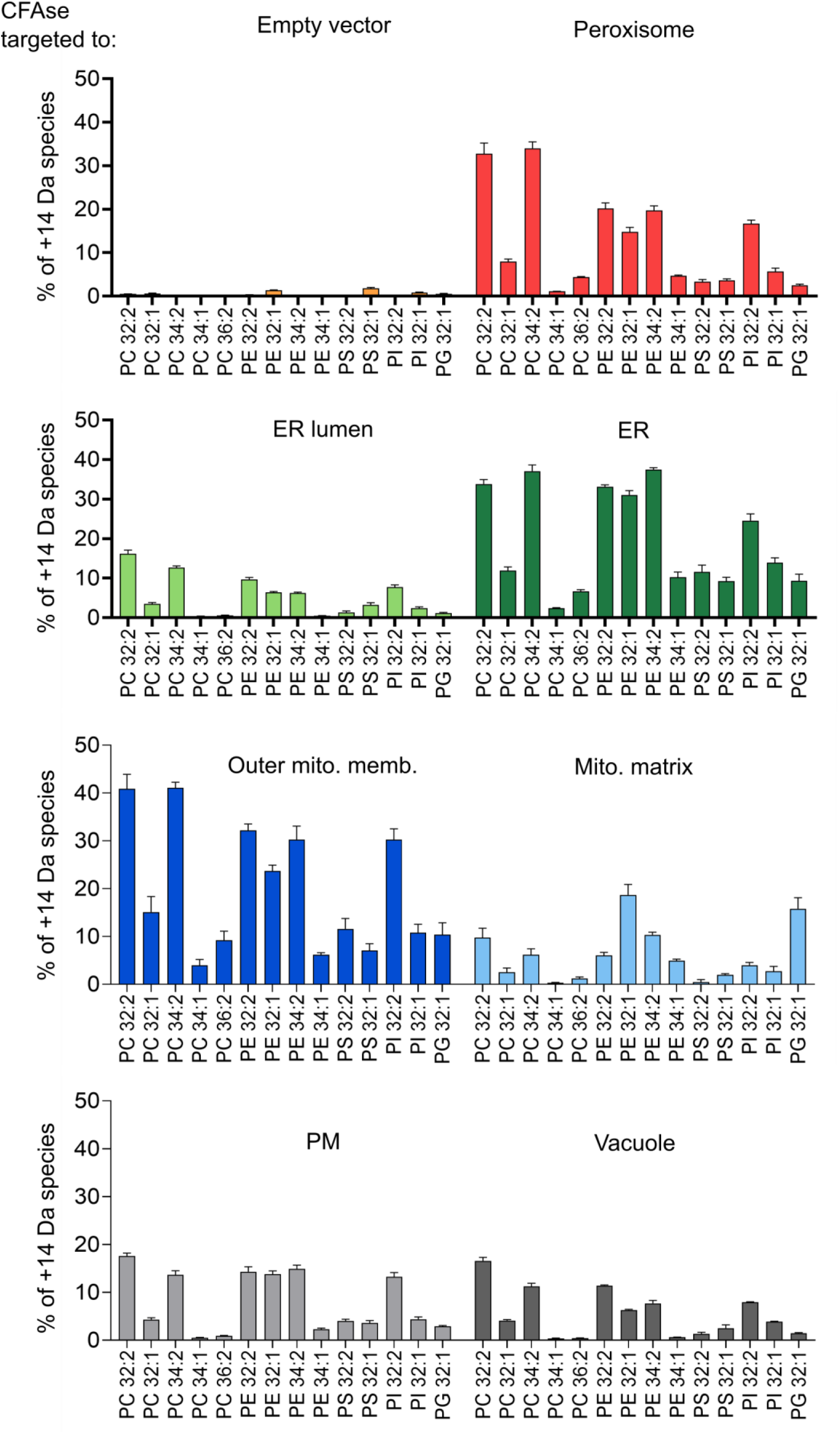
CFAse mass-tags a variety of phospholipid species in yeast upon targeting to different organelles. Bar plot showing the percentage of each indicated phospholipid species that is mass-tagged upon constitutive expression of organelle-targeted CFAse in yeast. Empty vector refers to a plasmid carrying no CFAse coding sequence.

##### Strategy to monitor ER-mitochondria lipid exchange *in vivo*

In order to monitor lipid transport between ER and mitochondria, we took advantage of the exclusive ER localization of the PE methyltransferases Cho2 and Opi3 as a way to introduce one of the two mass tags. We targeted CFAse to the mitochondrial matrix as a means to introduce the other mass tag. Since both CFAse and the methyltransferases use SAM as a methylene or methyl donor, respectively, we pulse-labeled cells with deuterated methionine (d-methionine) ^21^ and monitored the appearance of both singly- and doubly-labeled phospholipids, the latter being indicative of lipid transport between the ER and mitochondria (Fig 3A, B, C).

**Figure 3.**
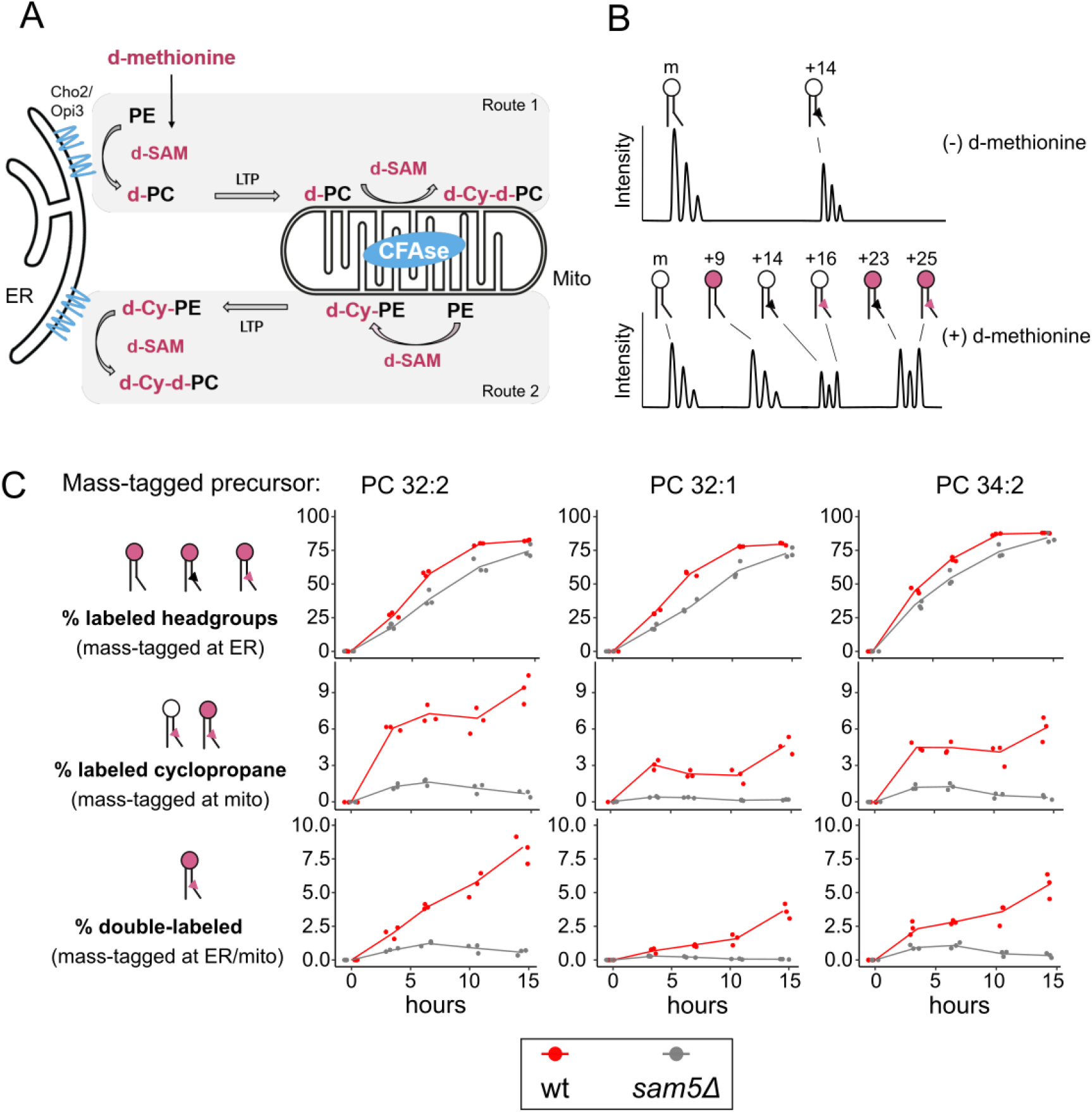
Monitoring ER-mitochondria lipid exchange using METALIC. *(A)* CFAse is targeted to the mitochondrial matrix, while the endogenous methytransferases, Cho2 and Opi3, localize to the ER. These enzymes at the two organelles serve to introduce distinct mass tags. Cells are pulse-labeled with deuterated methionine (d-methionine), resulting in deuterated SAM (d-SAM). In **Route 1**, the first mass-tagging occurs in ER, resulting in d-PC (+9 Da). When d-PC is transported by a lipid transport protein (LTP) to the mitochondrial matrix, a second mass tag (+16 Da) in the form of a deuterated cyclopropane group (d-Cy) is added by CFAse using d-SAM, resulting in d-Cy-d-PC (+25 Da). The doubly mass-tagged species can also result from **Route 2**, where PE in the mitochondria matrix can get a deuterated cyclopropane mass tag (+16 Da), which subsequently can be double mass-tagged (+9 Da) in the ER by the methyltransferases. *(B)* Theoretical mass spectra illustrating the different modifications of a precursor PC species. At steady state, in addition to the precursor, only the cyclopropylated +14 Da species (black triangle) is detected due to the constitutive expression of CFAse. Upon treatment with d-methionine, labeling of the headgroup at the ER (pink headgroups) results in the +9 Da shift. Headgroup labeling of the +14 Da species results in the detection of +23 Da species. Detection of +16 Da species indicates the labeling of the tail in PCs at mitochondria (red triangle). Double mass-labeling of both the headgroup (at ER) and the tail (at mitochondria) result in a +25 Da mass tag. *(C)* Line plot depicting the percentage of incorporation in the head group (sum of +9, +23 and +25 species), fatty acid tail (sum of the +16 and +25 species) and both (+25 species) after d-methionine pulse labeling of cells of the indicated genotype at the indicated time points. Three independent clones for each genotype were used.

As for the singly-labeled species, we monitor the +9 Da PC species resulting from the triple methylation of the headgroup by Cho2 and Opi3. We also monitor the +16 Da PC species, which results from cyclopropylation of PC species at mitochondria. The doubly labeled species has a +25 Da (9+16) mass shift, which results from triple methylation at the PC head group and fatty acid tail cyclopropylation (Fig 3B). In this arrangement, double mass-labeling can either result from the head group-labeled PC transported to mitochondria for the second mass-labeling (Fig 3A, Route 1) or cyclopropane-labeled PE transported from mitochondria to the ER for the second mass-labeling (Fig 3A, Route 2). Our measurements thus assess both transport directions between the two organelles.

Upon pulse labeling in wild-type cells, we observed a time-dependent increase in the fraction of lipids containing deuterated headgroups and deuterated cyclopropanes, among the most abundant PC species (i.e., 32:2, 32:1 and 34:2, Fig 3C, red line). While the incorporation of deuterated headgroups saturated close to 100%, the appearance of deuterated cyclopropanes in PCs saturated at much lower values, consistent with the fact that CFAse only modifies a fraction of the lipids (Fig 2). For all three major PC species, we observed the appearance of +25 Da double-labeled lipids in a time-dependent manner (Fig 3C, red line), indicative of ER-mitochondria lipid transport.

To experimentally validate the specificity of this METALIC strategy, we tested the dependency of mass tag-labeling on Sam5, the major transporter of SAM across the inner mitochondrial membrane ^22^. Since CFAse activity in the mitochondrial matrix is dependent on SAM, our prediction was that in the *sam5* mutant, mass-labeling at mitochondria should be severely impaired. Indeed, while the rate of headgroup labeling at ER was similar to wild-type cells (the small difference might be accounted for by a slower growth rate of *sam5Δ* mutants), we observed that the incorporation of deuterated cyclopropane rings, as well as double mass-labeling were severely reduced in *sam5Δ* mutant cells (Fig 3C, grey line). This result confirmed that CFAse targeting to the mitochondrial matrix was near complete with little activity outside of this compartment. Taken together, these results highlight the robustness and sensitivity of the approach for monitoring ER-mitochondria phospholipid exchange *in vivo*.

##### An auxin-inducible degron system to inactivate ERMES function

To clearly assess the redundant roles of ERMES and Vps13-Mcp1 complexes in lipid exchange, we sought to assay lipid transport upon co-inactivation of both pathways. As deletion of any one of the *ERMES* genes together with that of either *VPS13* or *MCP1* is synthetically lethal, we built an inducible system to acutely inactivate the ERMES complex. To this end, we C-terminally fused Mdm12 to an auxin-inducible degron (AID) ^23^ and expressed *At*TIR1, a plant auxin-dependent adapter for E3 ubiquitin ligases. FLAG-tagged Mdm12-AID was efficiently depleted after 1 hour of auxin treatment (Fig 4A). Moreover, ball-like mitochondria typical of *ermes* mutants were apparent upon auxin addition, in Mdm12-AID cells alone (Fig 4B) or in combination with *vps13Δ* or *mcp1Δ* (Fig 4D), as visualized microscopically in cells expressing a mitochondria matrix-targeted CFAse-mCherry construct. Finally, in the presence of auxin, cells expressing Mdm12-AID grew slower, and this defect was exacerbated by concomitant deletion of *VPS13* or *MCP1* (Fig 4C), in line with previous observations ^11,13,24^. Taken together, these results confirm that the auxin-dependent depletion of Mdm12 rapidly inactivates the ERMES complex, and thus allows testing its lipid transport activity *in vivo* using the METALIC assay.

**Figure 4.**
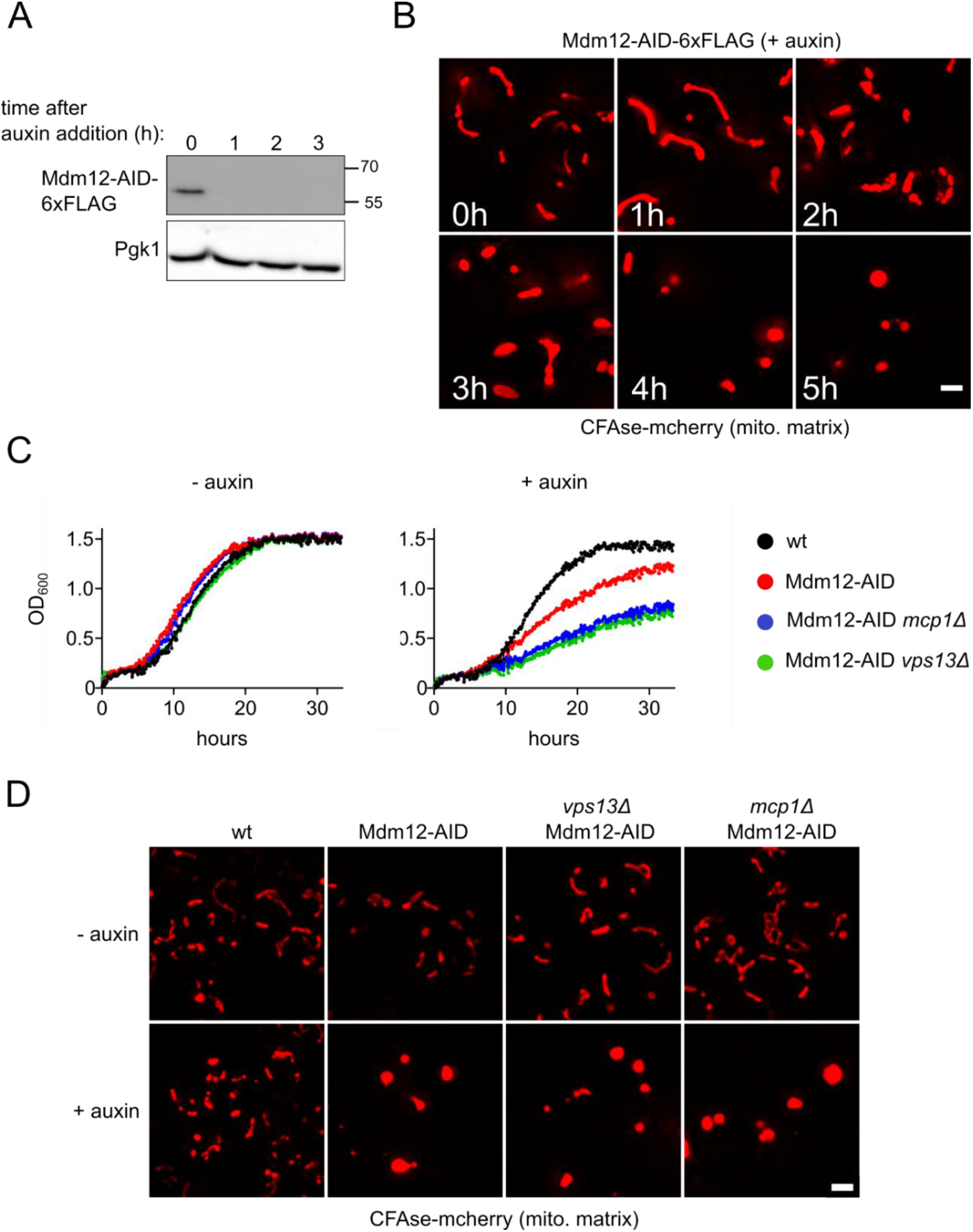
*(A)* An auxin-inducible degron (AID) system to acutely inactivate ERMES function in yeast. Cells co-expressing *At*TIR-9xMyc and Mdm12-AID-6xFLAG were treated with 0.5mM auxin, grown at 30°C and collected at defined time points (hours). Total protein extracts were analyzed by SDS-PAGE and Western blotting. Mdm12 was detected using a α-FLAG antibody. Phosphoglycerate kinase, which serves as a loading control, was detected using a α-PGK antibody. *(B)* Cells bearing the Mdm12 degron system and expressing the mitochondria matrix-targeted CFAse were treated with 0.5mM auxin, and imaged at the mentioned time points (hours). Images correspond to a maximum intensity projection of six Z-slices. *(C)* Growth of cells with the indicated genotypes was monitored using OD_600_ measurements in the absence or presence of 0.5mM auxin. *(D)* Localization of mCherry-tagged CFAse in the indicated strains, either in the absence of auxin or upon treatment with 0.5mM auxin for 7 hours.

##### The ERMES and Vps13-Mcp1 complexes both contribute to phospholipid exchange between ER and mitochondria *in vivo*

To unravel the roles of the ERMES and the Vps13-Mcp1 complexes in ER-mitochondria lipid exchange, we pulse-labeled cells with deuterated methionine and assayed mass tag-labeling upon inactivation of either one or both of the pathways (Fig 5A). The labeling kinetics of headgroups at the ER was similar in all the LTP mutants, indicating that despite different growth phenotypes, cells are metabolically active and generate newly synthesized PC headgroups at a comparable rate (Fig 5B, C, top panels).

**Figure 5.**
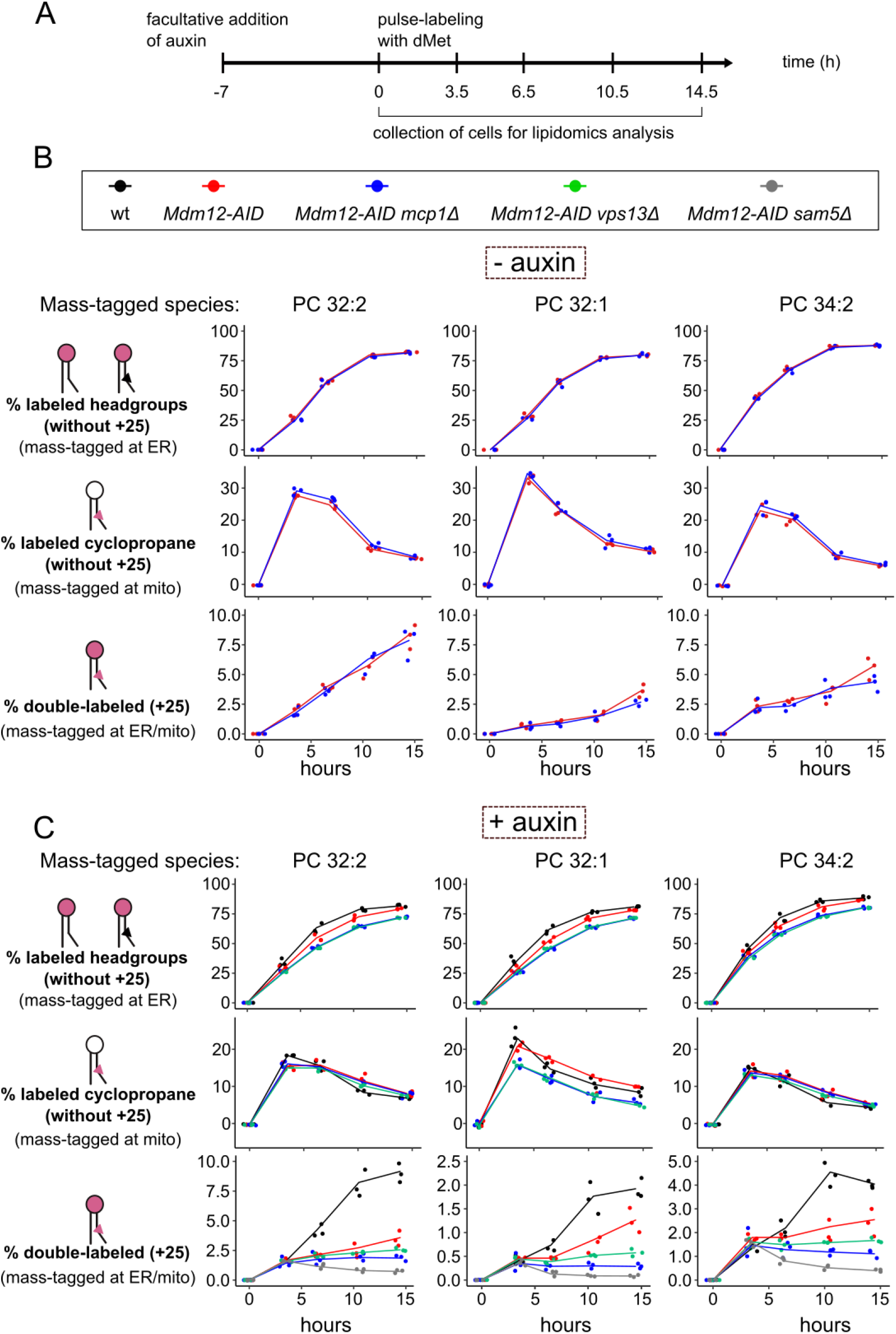
Kinetics of phospholipid exchange between ER and mitochondria. *(A)* Depiction of the time points at which cells were collected for lipidomics analysis upon pulse labeling with dMet, after treatment with 0.5mM auxin for 7 hours. *(B, C)* Line plots showing the fraction of the +9 Da, +16 Da and +25 Da species over time in the indicated genotypes either without *(B)* or with *(C)* auxin treatment. Three independent clones of each genotype were used.

To address the contribution of the Mcp1/Vps13 pathway alone to lipid transport, we assayed mass tag-labeling in *MDM-12-AID mcp1Δ* cells, in the absence of auxin (- auxin) to maintain ERMES function (Fig 5B). First, we quantified label incorporation at the headgroup and at the cyclopropane ring, independent of transport. To do so, we quantified all species with deuterated headgroup or cyclopropane rings with the exception of doubly mass-labeled species. While we observed a similar increase headgroup labeling, increased modification of the cyclopropane ring was transient, probably reflecting the fact that labeling at the headgroup is faster than at the cyclopropane ring, and thus by the end of the experiment most PC molecules bore deuterated headgroups. As shown in Fig 5B, *MCP1* deletion alone impacted neither headgroup nor cyclopropane ring labeling. We then quantified doubly mass-labeled lipids (+25 Da), indicative of ER-mitochondria lipid exchange. In this set-up, the abundance of doubly mass-labeled species increased monotonously in *MDM-12-AID* (surrogate wild-type) and *MDM-12-AID mcp1Δ* cells. While PC 32:2 was not affected, after 14.5 hours, there was a mild (25%) but significant reduction in the double mass labeling of PC 32:1 (*p*-value = 0.038) and a non-statistically significant reduction in PC 34:2 (*p*-value = 0.32) (Fig 5B, lower panels).

To assess the role of the ERMES complex, we treated cells bearing *MDM12-AID* with auxin for 7 hours before pulse-labeling (Fig 5A, Fig S1). While labeling of the headgroups and cyclopropane rings individually was unaffected upon Mdm12 depletion, the rate of double mass-labeling (+25 Da species) was reduced (Fig 5C, red lines). Interestingly, this reduction was more marked for the PC 32:2 species.

Finally, we assessed mass tagging in *MDM12-AID* cells with either *VPS13* or *MCP1* deleted. We observed a slight reduction in individual mass labeling of the headgroups and cyclopropane rings (at least for the PC32:1 species, Fig 5C, blue and green lines). Strikingly, however, double mass labeling was reduced close to background levels by co-inactivation of ERMES and either Vps13 or Mcp1. The effect of additional *MCP1* or *VPS13* deletion was more pronounced for PC 32:1 than for other PC species.

Thus, while both pathways might show some specificity with regard to the transported phospholipids, these results demonstrate that the ERMES and Vps13-Mcp1 complexes function in ER-mitochondria lipid exchange *in vivo*, providing a biochemical basis for their genetic redundancy.

## Discussion

Here we demonstrate the utility of METALIC as a novel strategy to probe interorganelle lipid transport *in vivo* using ER-mitochondria as a model organelle pair. Our unique survey of ER-mitochondria lipid transport, thus, ties hampering loose ends in our understanding of ER-mitochondria lipid exchange. Importantly, it demonstrates the contribution of two candidate pathways, for which direct *in vivo* evidence had been thus far missing or incomplete ^2–5,8,25^. We observe that Vps13-Mcp1 pathway contributes only minimally to ER-mitochondria lipid exchange. This weak phenotype of the Vps13-Mcp1 pathway correlates with the observation that neither *VPS13* nor *MCP1* deletion affect mitochondria morphology or yeast growth. On the other hand, the contribution of the ERMES complex to lipid transport is substantial, in line with the strong mitochondrial and growth phenotypes associated with *ermes* mutants. Moreover, the two pathways function in a redundant fashion, together accounting for the bulk of lipid transport between the two organelles, potentially explaining the synthetic lethality of mutants lacking both ERMES and Vps13-Mcp1 pathways.

Interestingly, the lipid transport defect observed upon ERMES inactivation is particularly striking for the doubly unsaturated species 32:2 and 34:2. By contrast, the 32:1 species was most affected in the *ermes vps13* or *ermes mcp1* double mutant cells. In fact, transport of this lipid was modestly but significantly affected by the loss of *MCP1* alone. Together, these findings suggest that lipid-binding pockets of LTPs, like Vps13 and ERMES components, could have preferences for specific fatty acids among phospholipids with the same headgroup.

The fact that CFAse can be directed to multiple cellular compartments makes it possible to study phospholipid transport between ER and any organelle of interest, as long as the targeting is stringent. In the case of mitochondrial matrix targeting, both microscopy results and the strong reduction of CFAse-mediated mass tagging in the *sam5*Δ mutant indicates that mistargeting of CFAse is negligible. In fact, we do not know if the residual activity of matrix-directed CFAse in the *sam5Δ* mutant is due to partial mistargeting of the enzyme or partial permeability of the mitochondrial membrane to SAM in the absence of Sam5.

Our data show that cyclopropane lipids can be synthesized and transported in yeast without noticeably perturbing cell function. Therefore, even if their behavior is likely not identical to unsaturated lipids, cyclopropane lipids can serve as useful tools within the METALIC approach to assay lipid transport, and unravel relative differences in different genetic backgrounds (lipid transport mutants) or physiological conditions. One limitation of the approach is that it does not inform on the directionality of transport (see Fig 3A, route 1 vs. route 2), nor whether the exchange is direct or involves intermediate compartments. Therefore, any rate calculated with METALIC cannot be used as a direct measure of lipid exchange. However, the effect of environmental and genetic perturbations on lipid traffic can be measured with METALIC, as we show here for ER-mitochondria lipid transport. One obvious caveat of enzyme-based methods like METALIC is that any perturbation can affect either lipid transport or the rate of mass tag incorporation. In METALIC, the rates of incorporation by both enzymes can be surveyed independently and used to normalize the rate of appearance of the doubly mass-tagged (and therefore transported) species.

The involvement of multiple redundant LTPs in interorganelle lipid transport appears to be the rule rather than the exception. Indeed, among the ~50 putative LTPs encoded in the yeast genome, none is truly essential for growth, indicating that redundant mechanisms can compensate for the absence of any one of them. Deciphering the contribution of these many LTPs, their redundancy, and their lipid preferences, therefore, constitutes an important challenge in cell biology. However, despite the central contribution of lipids to a plethora of cellular functions, our knowledge significantly lags behind that of DNA, RNA and proteins, to the point that we still ignore most of the routes taken by given lipids to travel from their synthesis site to their destination. This is because we do not have the equivalent tools (PCR, GFP-tagging) for lipids. The development of METALIC thus takes an important step forward and paves the way to elucidate the function of LTPs and lipid transport processes *in vivo*.

## Materials and methods

### Yeast strains and plasmids

Strains used in this study are listed in Table S1. Genomic integration of PCR fragments was done by homologous recombination ^26,27^. Gene deletions were confirmed by colony PCR. Plasmids and primers used in the study are listed in Tables S2 and S3, respectively. CFAse ORF was amplified from *E. coli*. A CFAse-mCherry construct was targeted to different organelles using the following targeting sequences; for the ER membrane, the C-terminal 20 residues of Ubc6 (230-250); for the ER lumen, the signal sequence of Kar2 (aa 1-41) at the N-terminus and a HDEL signal at the C-terminus; for the outer mitochondrial membrane, aa 1-30 of Tom70; for the mitochondrial matrix, aa 1-69 of subunit 9 of the F_0_-ATPase from *N.crassa*; for the peroxisome, full-length Pex3 at the N-terminus; for the plasma membrane, the PH domain of Osh1 (aa 268-388); for the vacuole, full-length Vac8 at the N-terminus.

### Microscopy

Cells were grown to mid-log phase in synthetic dextrose (SD)–uracil medium for selection of the mitochondrial matrix-targeted CFAse-mcherry plasmid. Images were acquired using a DeltaVision MPX microscope (Applied Precision) equipped with a 100× 1.40 NA oil UplanS-Apo objective lens (Olympus), a multicolor illumination light source, and a CoolSNAPHQ2 camera (Roper Scientific). Image acquisition was done at room temperature. Images were deconvolved with SoftWoRx software using the manufacturer’s parameters. Images were processed further using FIJI ImageJ bundle.

### Protein extraction and western blotting

1 OD_600_ of mid-log phase cells were collected by centrifugation and precipitated using 10% trichloroacetic acid for 20 min at 4°C. After centrifugation at 13,000 *g* for 5 min, pellets were washed with ice-cold acetone. Pellets were air-dried and resuspended in 30 μl of 1× SDS sample buffer (60 mM Tris, pH 6.8, 2% SDS, 10% glycerol, 5% 2-mercaptoethanol, and 0.005% bromophenol blue), and boiled for 3 min. Samples were resolved on a 12% SDS-PAGE gel, and after transfer on a PVDF membrane, proteins were detected using α-FLAG (Sigma Aldrich) or α-PGK antibody.

### Pulse-labeling, lipid extraction and mass spectrometry analysis

Pre-cultures in synthetic minimal medium were diluted to 0.05 OD_600_/ml in 25 ml and treated with 0.5 mM auxin for 7 hours. Next, cells were pulse labeled with 2 mM deuterated methionine and grown at 30 °C. At the indicated time points, 8 OD_600_ of cells was pelleted, snap-frozen and stored at −80°C. Lipids were extracted as described previously with minor modifications ^28^. Briefly, cells were washed in ice-cold water and subsequently resuspended in 1.5 ml of extraction solvent containing ethanol, water, diethyl ether, pyridine, and 4.2 N ammonium hydroxide (v/v 15:15:5:1:0.18). After the addition of 300μL glass beads, samples were vortexed vigorously for 5 minutes and incubated at 60 °C for 20 min. Cell debris were pelleted by centrifugation at 1,800 ×g for 10 min and the supernatant was dried under a stream of nitrogen. The dried extract was resuspended in 1 ml of water-saturated butanol and sonicated for 5 min in a water bath sonicator. 500 μl of water was added and vortexed further for 2 min. After centrifugation at 3000 ×g, the upper butanol phase was collected, dried under a stream of nitrogen and resuspended in 50% methanol for lipidomics analysis.

LC analysis was performed as described previously with several modifications ^29^. Phospholipids were separated on a nanoAcquity UPLC (Waters) equipped with a HSS T3 capillary column (150 m x30mm, 1.8 m particle size, Waters), applying a 10 min linear gradient of buffer A (5 mM ammonium acetate in acetonitrile/water 60:40) and B (5 mM ammonium acetate in isopropanol/acetonitrile 90:10) from 10% B to 100% B. Conditions were kept at 100% B for the next 7 min, followed by a 8 min re-equilibration to 10% B. The injection volume was 1 μL. The flow rate was constant at 2.5 μl/min.

The UPLC was coupled to QExactive mass spectrometer (Thermo) by a nanoESI source (New Objective Digital PicoView® 550). The source was operated with a spray voltage of 2.9 kV in positive mode and 2.5 kV in negative mode. Sheath gas flow rate was set to 25 and 20 for positive and negative mode, respectively. MS data was acquired using either positive or negative polarization, alternating between full MS and all ion fragmentation (AIF) scans. Full scan MS spectra were acquired in profile mode from 107-1600 m/z with an automatic gain control target of 1e6, an Orbitrap resolution of 70,000, and a maximum injection time of 200 ms. AIF spectra were acquired from 107-1600 m/z with an automatic gain control value of 5e4, a resolution of 17,500, a maximum injection time of 50 ms and fragmented with a normalized collision energy of 20, 30 and 40 (arbitrary units). Generated fragment ions were scanned in the linear trap. Positive-ion-mode was employed for monitoring PC and negative-ion-mode was used for monitoring PS and PE. Lipid species were identified based on their m/z and elution time. We used a standard mixture comprising PS 10:0/10:0, PE 17:0/17:0, PC 17:0/17:0, PG 17:0/17:0 and PI 12:0/13:0 for deriving an estimate of specific elution times. Lipid intensities were quantified using the Skyline software ^30^. For each phospholipid, signal was integrated for the precursor species (m), cyclopropane species (m_+14_) and species that appear upon pulse-labeling with deuterated methionine (m_+9_, m_+16_, m_+23_, m_+25_). Fraction of cyclopropylated species (Fig 2) upon constitutive expression of CFAse was calculated as (m_+14_) / (m + m_+14_). Fraction of labeled headgroups (Fig 3), was calculated as (m_+9_ + m_+23_ + m_+25_) / (m + m_+9_ + m_+14_ + m_+16_ + m_+23_ + m_+25_). Fraction of labeled cyclopropane (Fig 3) was calculated as (m_+16_ + m_+25_) / (m + m_+9_ + m_+14_ + m_+16_ + m_+23_ + m_+25_). Fraction of labeled headgroups and cyclopropane, independent of transport (Fig 5), were calculated as (m_+9_ + m_+23_) / (m + m_+9_ + m_+14_ + m_+16_ + m_+23_ + m_+25_) and (m_+16_) / (m_+14_ + m_+16_ + m_+23_ + m_+25_), respectively. Fraction of doubly labeled mass-tagged species (Fig 3 and 5) was calculated as (m_+25_) / (m + m_+9_ + m_+14_ + m_+16_ + m_+23_ + m_+25_).

## Author contributions

A.T. John Peter and B. Kornmann conceived the study. A.T. John Peter designed and performed the experiments with the supervision of M. Peter and B. Kornmann. A.T. John Peter and B. Kornmann analyzed data. A.T. John Peter and B. Kornmann wrote the manuscript with input from M. Peter.

## Acknowledgements

We are thankful to members of the Peter and Kornmann laboratories for insightful discussions and helpful suggestions. Microscopy analysis was carried out at the ETH Zürich ScopeM facility, and we thank Dr. Tobias Schwarz for outstanding technical support. Lipidomics measurements were performed at the Functional Genomics Center Zurich (FGCZ). We especially thank Dr. Sebastian Streb and Dr. Endre Laczko of the FGCZ Metabolomics division for establishing and optimizing lipidomics workflows, and for excellent technical guidance. Ylp204-pADH1-*At*TIR1-9myc was a gift from Helle Ulrich (Addgene plasmid #99532), and we are grateful to Dr. Philipp Kimmig for providing us the *At*TIR1 construct in a pRS303 backbone.

The Kornmann lab is supported by grants from the Swiss National Science Foundation (SNSF, 31003A_179549) and the Wellcome Trust (214291/A/18/Z). Work in the Peter laboratory was supported by the SNSF, the Synapsis Foundation and ETH Zürich. A.T.J.P was supported by ETH Zürich/Institute of Biochemistry and a Spark grant of the SNSF (CRSK-3_190364). The authors declare no competing financial interests.

## Supplementary Figure

**Figure S1.**
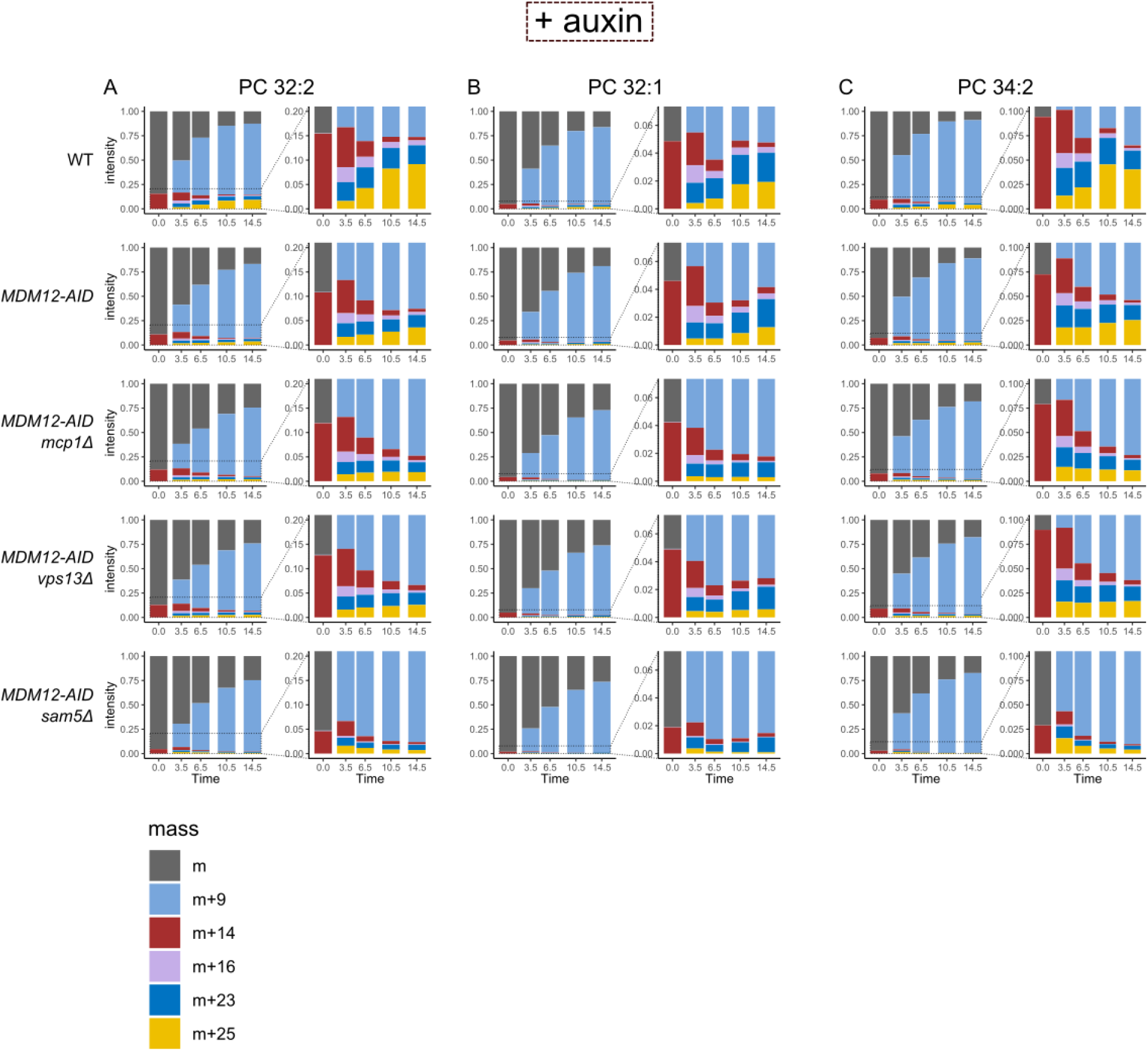
Normalized bar plot showing the relative changes in intensities of different labeled species with time (hours), from the onset of d-methionine labeling in auxin-treated cells. The changes are shown for three different phospholipid species, PC 32:2 (*A*), PC 32:1 (*B*) and PC 34:2 (*C*). Inset shows magnified views to better follow the intensity changes in species with a cyclopropane ring.

## Supplementary Tables

**Table S1.**
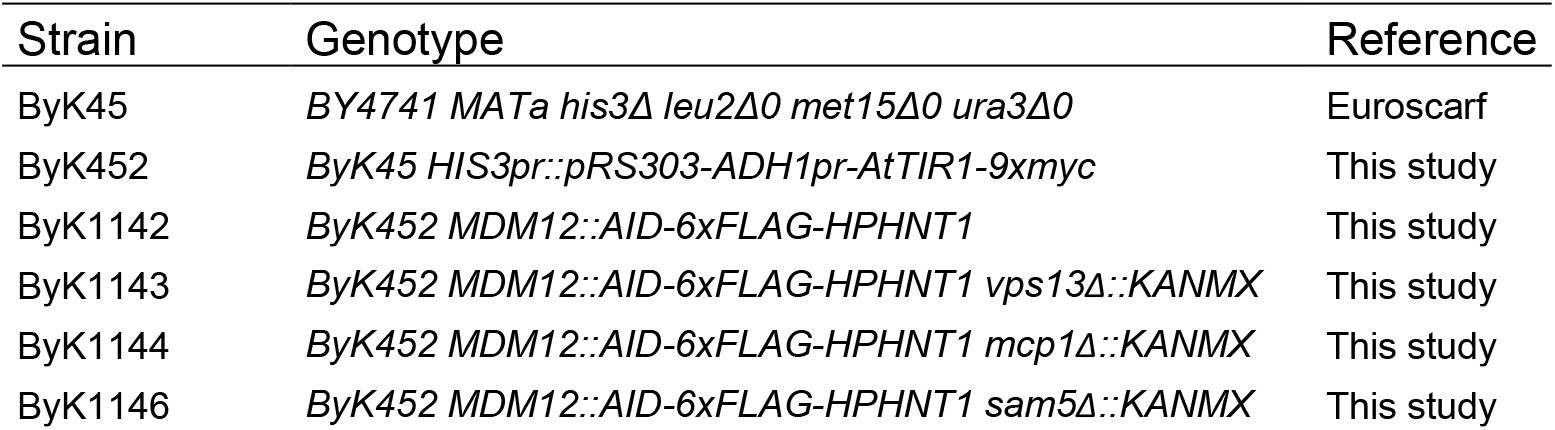
Yeast strains used in this study.

**Table S2.**
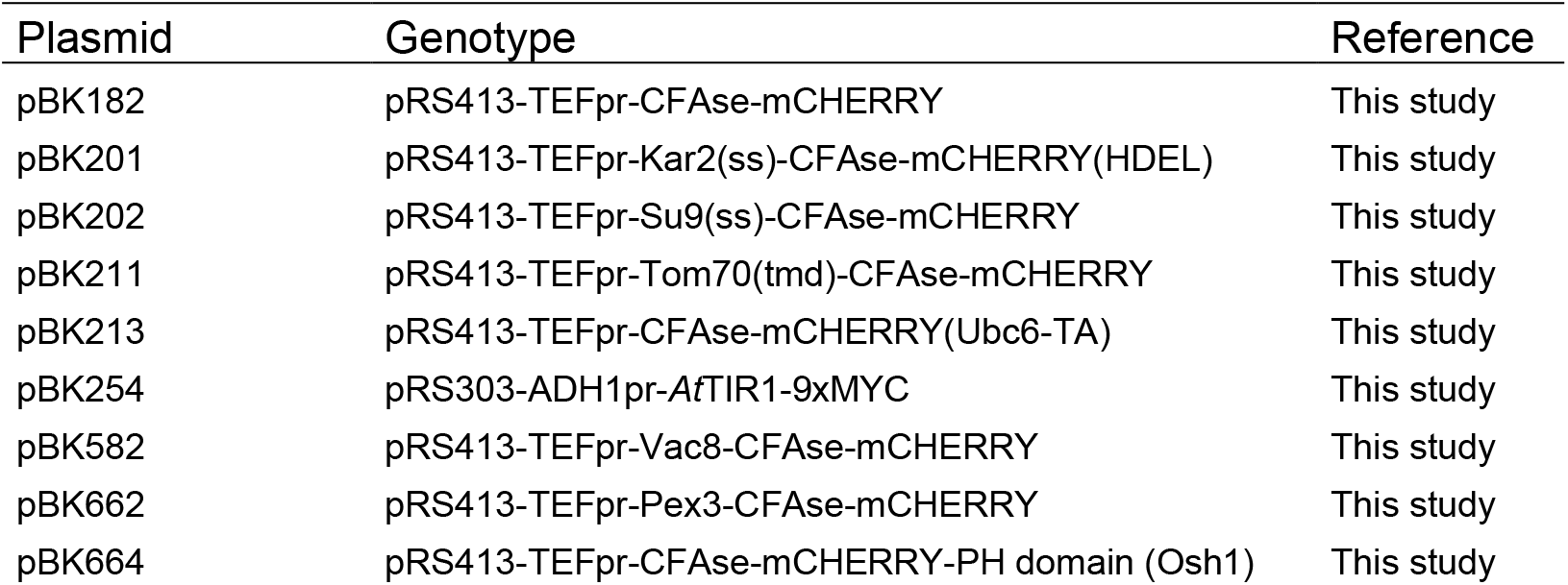
Plasmids used in this study.

**Table S3.**
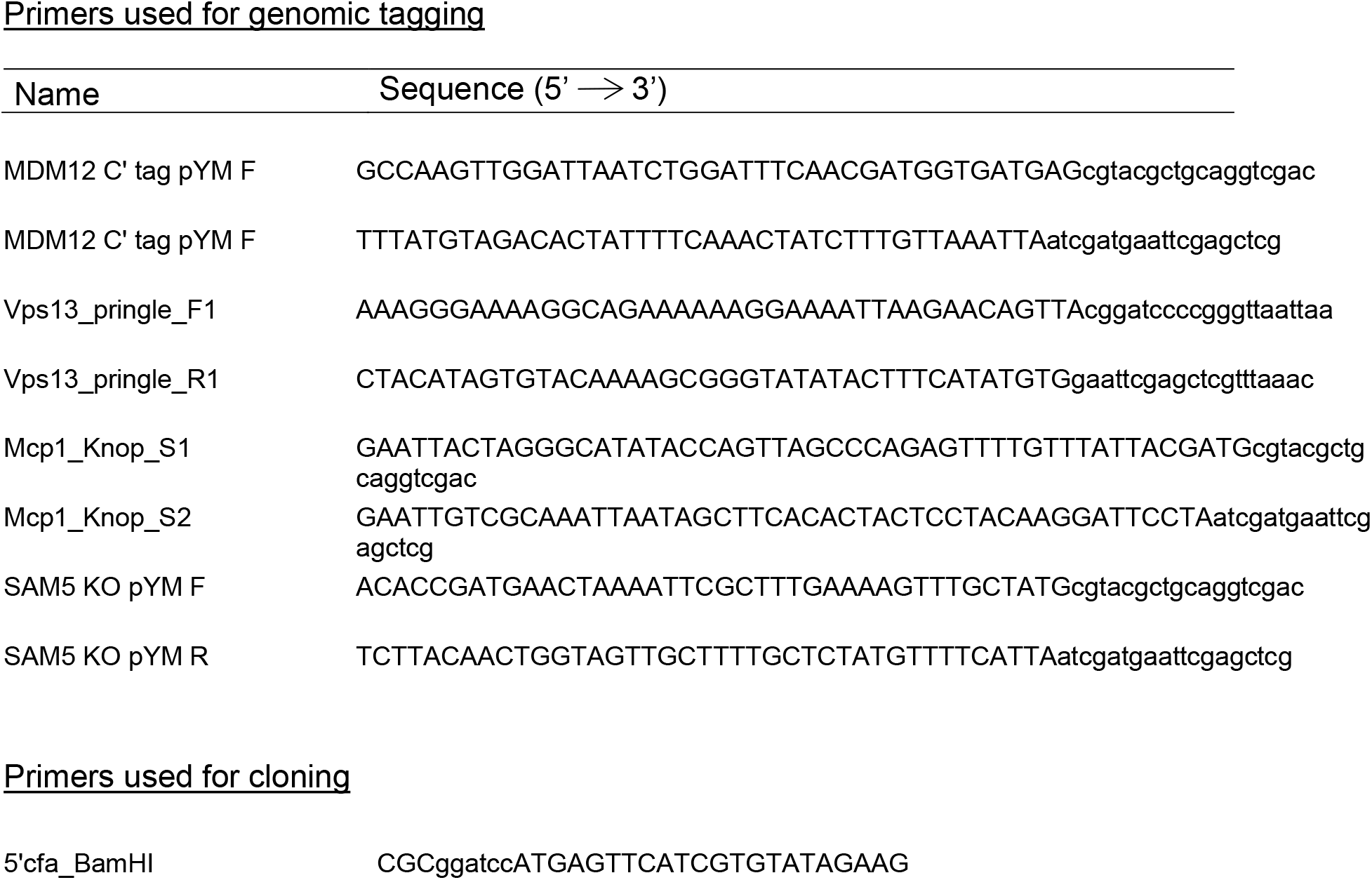
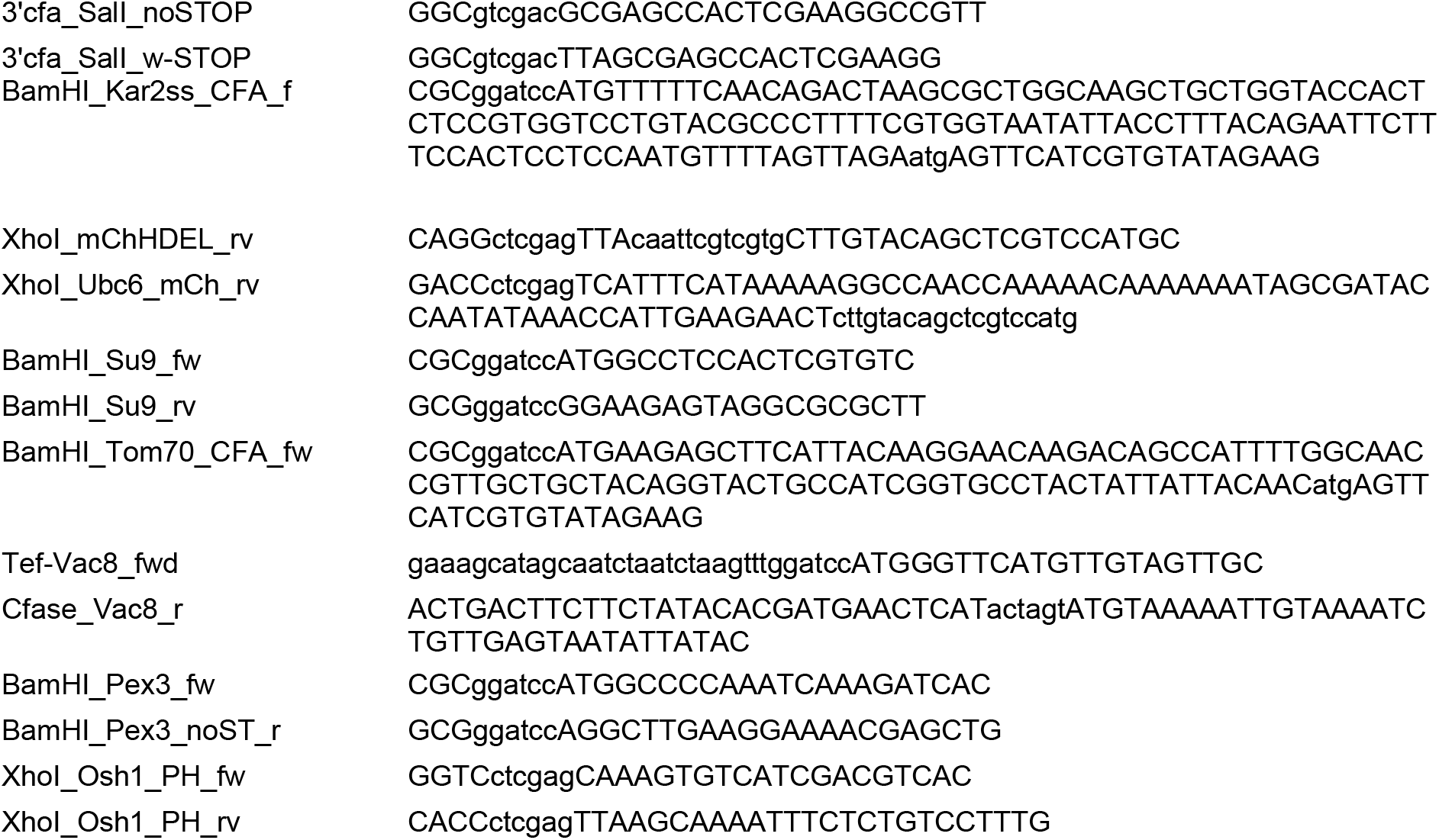
Primers used in this study.

